# Hidden Players: Latent Viruses Can Modulate Plant Disease Outcomes in Coinfection Contexts

**DOI:** 10.64898/2026.10.21.760619

**Authors:** Guillaume Lafforgue, Santiago F. Elena

## Abstract

Latent viruses are ubiquitous components of plant viromes and are likely the rule rather than the exception in natural host populations. Yet, despite their prevalence, little is known about how they influence the outcome of infections by acute viruses. Here, we used a factorial design to test the effects of two latent secoviruses, apple latent spherical virus (ALSV) and Arabidopsis latent virus 1 (ArLV1), on infections by the acute viruses tobacco rattle virus (TRV) and turnip mosaic virus (TuMV). Coinfections were evaluated across eight *Arabidopsis thaliana* genotypes, including wild-type and mutants impaired in RNA silencing, DNA methylation and hormone-signaling pathways. Latent-virus effects on both viral accumulation and disease severity were highly dependent on the combination of host and viral genotypes. Hormonal pathways played a prominent role in mediating these interactions, with recurrent responses involving jasmonic acid, salicylic acid and ethylene signaling. ALSV generally attenuated disease severity and acute-virus accumulation, whereas ArLV1 produced both antagonistic and synergistic outcomes depending on host background. Notably, ArLV1 coinfection increased TuMV-induced disease severity in some genotype, an effect never observed with ALSV. These results demonstrate that latent viruses can substantially modify disease expression and viral fitness and highlight the need to account for the pre-existing virome when predicting plant disease outcomes, viral emergence, and epidemiological dynamics.

## 1. Introduction

For more than a century, plant virology has largely been framed by a reductionist paradigm in which disease results from the interaction between a single virus and a single host. This perspective emerged from the study of major epidemics in agricultural monocultures (Elena et al., 2014; Roossinck, 2015). However, the development of virus ecology and the widespread adoption of high-throughput sequencing have revealed a much more complex reality. In natural and agricultural ecosystems, plants are frequently infected by multiple viruses simultaneously, forming dynamic viral communities or viromes (Mascia and Gallitelli, 2016; McLeish et al., 2019; Alcaide et al., 2020b).

Mixed infections are now recognized as the rule rather than the exception. Metagenomic surveys indicate that 60 - 70% of wild plants harbor more than one virus, often including phylogenetically distant species occupying the same host tissues (Roossinck, 2015; Lefeuvre et al., 2019). Such coinfections are promoted by polyphagous vectors, vegetative propagation and intensive agricultural practices that facilitate the maintenance and transmission of multiviral complexes (McLeish et al., 2019; Alcaide et al., 2020a). Consequently, infection outcomes cannot be understood solely from the properties of individual viruses. Virus-virus interactions range from antagonistic relationships, such as competition and cross-protection, to synergistic interactions that increase viral accumulation and disease severity (Syller, 2012; Syller and Grupa, 2015). These interactions can profoundly affect viral fitness, tissue tropism, transmission, and evolutionary trajectories (Mascia and Gallitelli, 2016).

An important but historically overlooked component of plant viromes is the large group of latent or asymptomatic viruses. Metagenomic studies have shown that many viruses persist in plants without causing visible symptoms or detectable fitness costs and may even provide benefits such as improved tolerance to abiotic stress or protection against other pathogens (Roossinck, 2015; Khankhum and Valverde, 2018; Fukuhara et al., 2020). Yet, despite their prevalence, the ecological and evolutionary consequences of latent viruses remain poorly understood. In particular, it is unclear whether latent viruses modify host susceptibility, alter immune responses, or influence the outcome of subsequent infections by acute viruses. Resolving these questions is essential for understanding viral emergence and disease dynamics in natural and agricultural ecosystems (Lefeuvre et al., 2019; Alcaide et al., 2020b).

Evidence from several pathosystems suggests that pre-existing viral infections can substantially alter disease outcomes. Cross-protection, whereby a prior infection limits subsequent infection by a related virus, is one well-known example (Paudel and Sanfaçon, 2018). More broadly, latent viruses can influence viral population dynamics, symptom severity and transmission processes. For instance, interactions between latent and acute variants of grapevine Pinot gris virus modulate disease expression in grapevines (Bertazzon et al., 2025). Similarly, latent infection by pepper cryptic virus 1 (PVC-1) deters aphids, unlike cucumber mosaic virus (CMV), which attracts them to facilitate its spread. This contrast suggests that PVC-1 may establish a beneficial relationship with its host plant by reducing both aphid herbivory and the transmission of acute viruses (Safari et al., 2019; Takahashi et al., 2019). These observations suggest that latent viruses may act as hidden modulators of acute virulent infections.

To investigate this possibility, we used two latent viruses that establish asymptomatic infections in *Arabidopsis thaliana*: apple latent spherical virus (ALSV; species *Cheravirus mali*, genus *Cheravirus*, family *Secoviridae*) and Arabidopsis latent virus 1 (ArLV1; species *Comovirus arabidopsis*, genus *Comovirus*, family *Secoviridae*). ALSV can infect *A. thaliana* asymptomatically and persist through meristem colonization and seed transmission (Igarashi et al., 2009). ArLV1 is frequently detected in *A. thaliana* populations and in publicly available RNA-seq datasets, indicating widespread and often unnoticed infections (Verhoeven et al., 2023). Although both viruses are latent and belong to the same family, ArLV1 is naturally adapted to *A. thaliana*, whereas ALSV is not. This contrast provides an opportunity to determine whether the effects of latent infections depend on host adaptation.

As models of acute infections, we selected tobacco rattle virus (TRV; species *Tobravirus tabacci*, genus *Tobravirus*, family *Virgaviridae*) and turnip mosaic virus (TuMV; species *Potyvirus rapae*, genus *Potyvirus*, family *Potyviridae*). TuMV is a highly adapted and pathogenic virus of *Brassicaceae* that induces severe symptoms in the *A. thaliana* Col-0 accession (Martín et al., 1999; González et al., 2019; Corrêa et al., 2020; Navarro et al., 2022; Ambrós et al., 2024). In contrast, TRV possesses a distinct biology, including a relatively weak viral suppressor of RNA (VSR) silencing compared with the potent HC-Pro suppressor of TuMV (Martínez-Priego et al., 2008). Together, these viruses provide complementary systems for assessing whether the effects of latent-virus coinfection are virus-specific or represent more general phenomena.

Because virus-virus interactions are likely mediated by host defense mechanisms, we examined coinfections across a panel of *A. thaliana* mutants impaired in major antiviral pathways. These mutants target RNA silencing (double mutant *dcl2 dcl4*), salicylic acid (SA) production (*sid2*) and signaling (*npr1*), jasmonic acid (JA) signaling (*jin1*), ethylene (ET) signaling (*ein2*), and genome-wide epigenetic regulation (*ddm1* and double mutant *drm1 drm2*). Previous work has shown that disruption of these pathways modifies susceptibility to viral infection and can alter the selective environment experienced by TuMV (Navarro et al., 2022; Ambrós et al., 2024; Olmo-Uceda et al., 2024; Elena et al., 2025; Elena 2026).

Despite growing recognition of mixed infections, the mechanisms governing interactions between latent and acute viruses remain poorly understood. In particular, we do not know whether latent viruses alter the accumulation and pathogenicity of secondary acute infections, nor whether these effects depend on specific host defense pathways. To address these questions, we employed a factorial design combining two latent viruses (ALSV and ArLV1), two acute viruses (TRV and TuMV) and a panel of *A. thaliana* defense mutants. This framework allows us to determine how latent-virus coinfection influences acute infections and to identify the host mechanisms that mediate these interactions.

## 2. Materials and methods

### 2.1 Plants, viruses, and growth conditions

A collection of eight different *A. thaliana* (L.) HEYNH genotypes of the Col-0 accession were used for this study (Table 1). This set covers a number of single or double mutants affecting different genes involved in the main antiviral defense pathways: (*i*) the wild-type Col-0 accession, the following simple and double mutants all in Col-0 background: (*ii*) *dcl2 dcl4* (RNA silencing), (*iii*) *drm1 drm2* (DNA methylation), (*iv*) *ddm1* (chromatin remodeling/methylation), (*v*) *npr1-1* (SA signaling), (*vi*) *sid2-1* (SA biosynthesis), (*vii*) *ein2-1* (ET signaling), and (*viii*) *jin1* (JA signaling).

**TABLE 1.**
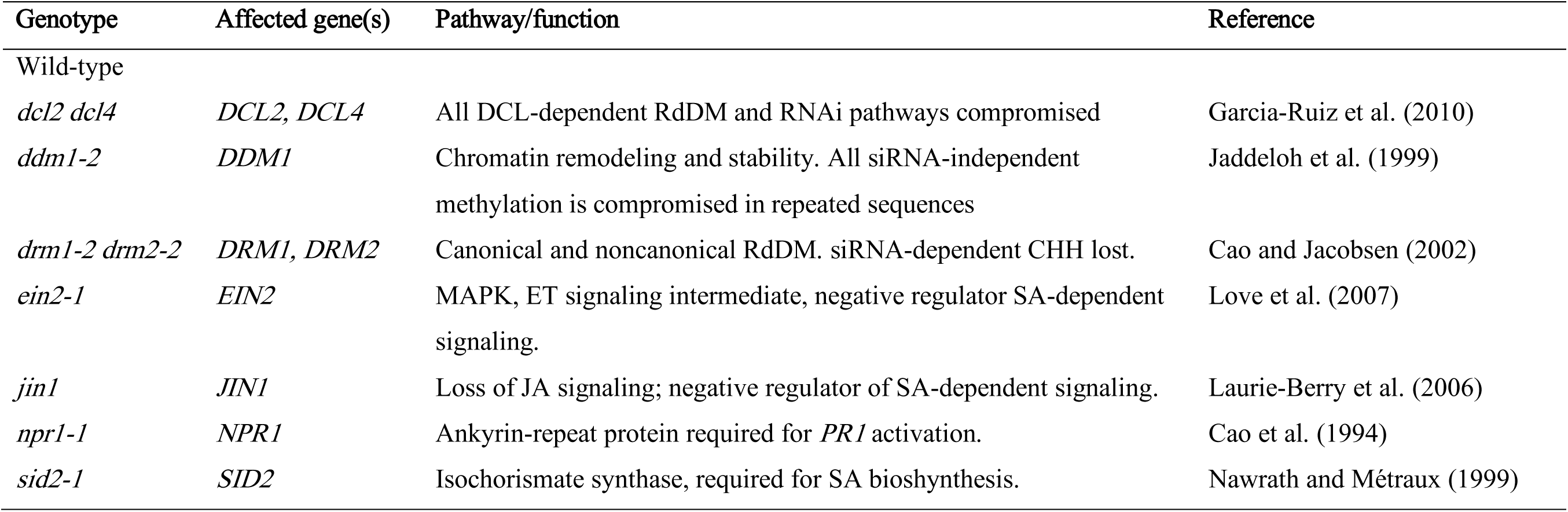
***Arabidopsis thaliana* genotypes used in this study.**

| Genotype | Affected gene(s) | Pathway/function | Reference |
| --- | --- | --- | --- |
| Wild-type |  |  |  |
| *dcl2 dcl4* | *DCL2, DCL4* | All DCL-dependent RdDM and RNAi pathways compromised | Garcia-Ruiz et al. (2010) |
| *ddm1-2* | *DDM1* | Chromatin remodeling and stability. All siRNA-independent methylation is compromised in repeated sequences | Jaddeloh et al. (1999) |
| *drm1-2 drm2-2* | *DRM1, DRM2* | Canonical and noncanonical RdDM. siRNA-dependent CHH lost. | Cao and Jacobsen (2002) |
| *ein2-1* | *EIN2* | MAPK, ET signaling intermediate, negative regulator SA-dependent signaling. | Love et al. (2007) |
| *jin1* | *JIN1* | Loss of JA signaling; negative regulator of SA-dependent signaling. | Laurie-Berry et al. (2006) |
| *npr1-1* | *NPR1* | Ankyrin-repeat protein required for *PR1* activation. | Cao et al. (1994) |
| *sid2-1* | *SID2* | Isochorismate synthase, required for SA biosynthesis. | Nawrath and Métraux (1999) |

In all experiments, plants were maintained in a BSL2 climatic chamber under a photoperiod of 8 h light (LED tubes at PAR 90 - 100 μmol/m^2^/s) at 24 °C and 16 h dark at 20 °C and 40% relative humidity in a mixture of 50% DSM WNR1 R73454 substrate (Kekkilä Professional, Vantaa, Finland), 25% grade 3 vermiculite and 25% 3–6 mm perlite. Pest management was performed by the introduction of *Stratiolaelaps scimitus* and *Steinernema feltiae* (Koppert Co., Málaga, Spain).

TRV was delivered from the infectious plasmids pTRV1 (GenBank accession AF406990) and pTRV2 (GenBank accession AF530055) (Liu et al., 2002).

TuMV-DV was obtained after an evolution experiment performed in *A. thaliana* Col-0 (González et al., 2019) that began with the ancestral TuMV YC5 strain from calla lily (*Zantedeschia* sp.) cloned in the infectious plasmid p35STunos (GenBank accession AF530055.2) (Chen et al., 2003). TuMV-DV shows strong symptoms severity and faster disease progression in *A. thaliana* Col-0 (González et al., 2019; Corrêa et al., 2020).

ALSV was propagated from the infectious clones pBICAL1 (GenBank accession NC_003787) and pBICAL12 (GenBank accession NC_003788) described by Li et al. (2004).

ArLV1 was kindly provided by Dr. Rene A.A. van der Vlugt as infected Col-0 *A. thaliana* material (Verhoeven et al., 2023).

### 2.2 Virus inoculation

ALSV and TRV inocula were produced in *Nicotiana benthamiana* DOMIN: leaves were rub-inoculated (ALSV) or agroinfiltrated (TRV), and infected tissue was collected at 12 dpi, frozen, and ground to a powder stock. TuMV infectious sap was obtained from infected *A. thaliana* stock plants (González et al., 2019). ArLV1 stock was produced by rub-inoculating *N. benthamiana* plants with sap from the ArLV1-infected Col-0 original tissue.

For inoculations, the viral powder was first quantified by absolute reverse transcription real-time quantitative polymerase chain reaction (RT-qPCR; see paragraph 2.4) to ensure a concentration of 3×10^12^ RNA molecules/μL. The quantified material was then diluted in inoculation buffer (50 mM phosphate buffer, pH 7.0, 3% PEG6000, and 10% Carborundum).

For single infections, each virus (SAP1 or SAP2) was separately mixed with the inoculation buffer at 1:1 (v/v) ratio. For coinfection, SAP1 and SAP2 were combined in equal proportions.

Five μL of the inoculum was gently rubbed onto two leaves per plant. Plants were all inoculated when they reached growth stage 3.5 in the Boyes et al. (2001) scale. This synchronization ensures that all hosts were at the same phenological stage when inoculated.

Twelve plants per Arabidopsis genotype × Infection type combination were inoculated in each experimental repetition. The TRV-related experiments (TRV, TRV/ALSV and TRV/ArLV1) were repeated twice in independent trials, while the TuMV-related experiments (TuMV, TuMV/ALSV and TuMV/ArLV1) were repeated three times.

### 2.3 Phenotyping infection

Two response variables were scored. (*i*) Disease severity was evaluated daily from 4 to 14 days post-inoculation (dpi) on a semiquantitative discrete scale ranging from 0 for no infection or asymptomatic infection to 5 for plants showing generalized necrosis and wilting (Corrêa et al., 2020; Butković et al., 2021). (*ii*) Viral load, measured by RT-qPCR as the number of viral genomes per ng of total RNA in the plants, as described below.

### 2.4 Total RNA extractions and quantification of viral load

At 14 dpi, plants were separated by the presence of symptoms. Symptomatic and non-symptomatic plants were pooled into three groups and frozen with liquid N_2_. This frozen plant tissue was homogenized into a fine powder using a Mixer Mill MM400 (Retsch GmbH, Haan, Germany).

Next, aliquots of approximately 0.1 g of ground tissues were used for total RNA (RNAt) extractions. The RNAt was extracted with the Agilent Plant RNA isolation Mini kit (Agilent Technologies, Santa Clara CA, USA). Three aliquots of RNAt per each sample were separated and their concentration adjusted at approximately 10 ng/μL to estimate viral load. Viral load was quantified in each of the three aliquots by RT-qPCR using standard curves and the pairs of primers of TRV, TuMV, ALSV, and ArLV1 shown in Table 2.

**TABLE 2.**
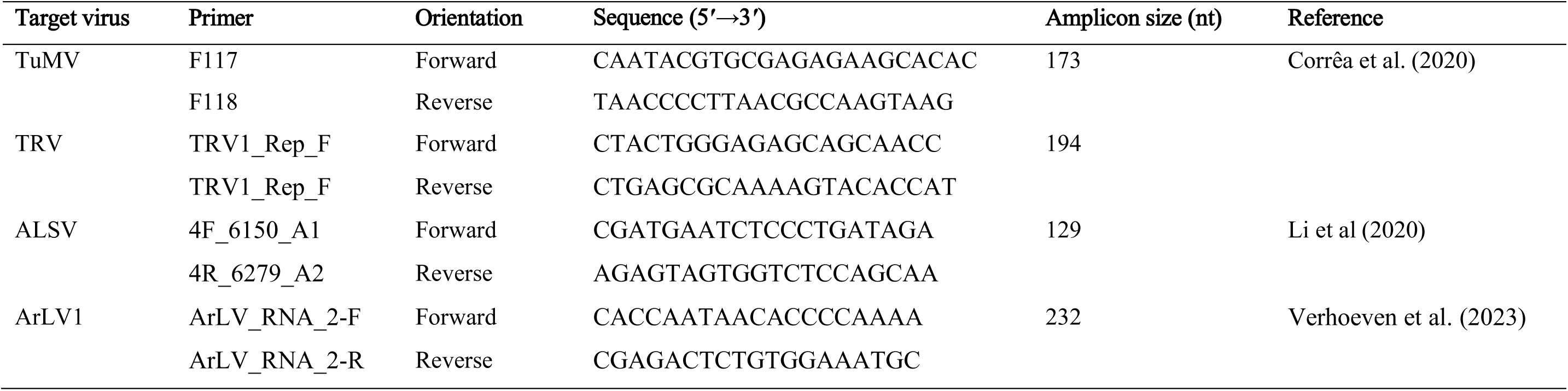
Primers used for RT-qPCR quantification of viral loads.

Briefly, standard curves were constructed using five serial dilutions of *in vitro* transcribed RNAs from the corresponding infectious clone. In the case of TRV, RNA1 was used chosen. Dilutions were made in sap generated from noninfected plants. Amplification reactions were run in a 20 μL volume using the qPCRBIO SyGreen^®^ 1-Step kit (PCR Biosystems Inc., East Hartford CT, USA). The cycling conditions consisted of an RT phase of 5 min at 42 °C and 10 min at 95 °C followed by a PCR stage consisting of 40 cycles of 5 s at 95 °C and 34 s at 60 °C, and a final melt curve profile consisting of 15 s at 95 °C, 1 min at 60 °C and 15 s at 95 °C. Negative controls consisted of healthy RNAt plant extract (mock-inoculated noninfected control) and water. For each sample, three technical replicates were quantified. The results were analyzed using the StepOne software 2.2.2 (Applied Biosystems, Foster City CA, USA).

### 2.5 Statistical analyses

All statistical analyses were performed in R version 4.6.0 under RStudio version 2026.08.2+200 (Posit Software PBC, Boston MA, USA) using Bayesian hierarchical models fitted with the ‘brms’ package version 2.23.0 (Bürkner, 2017), which interfaces with Stan for Hamiltonian Monte Carlo (HMC) sampling. Two modeling frameworks were employed depending on the response variable. For viral load (log_10_ viral genomes per ng of total RNA), technical replicates were averaged per biological replicate. Bayesian linear mixed models (BLMMs) were fitted separately for each quantified virus (TRV, TuMV and ArLV1) using a Gaussian likelihood. The fixed-effects structure included Arabidopsis genotype × Infection type, with experimental block and biological replicate nested as random intercepts (1 | Block/Biological replicate). The variable Infection type takes values 1 (single infection with the corresponding virus), 2 (double infection with first alternative virus) and 3 (double infection with second alternative virus). Biorep corresponds to 3 pools of 4 plants.

For disease severity progression, Bayesian ordinal generalized linear mixed models (BGLMMs) were fitted using a cumulative distribution with a logit link function, appropriate for the ordered categorical symptom scale (0 - 5). Separate models were fitted for the TRV pathosystem (comprising TRV single infection, TRV/ArLV1 and TRV/ALSV coinfections) and the TuMV pathosystem (comprising TuMV single infection, TuMV/ArLV1 and TuMV/ALSV coinfections). The fixed-effects structure included Arabidopsis genotype × Infection type × dpi, with experimental batch and plant identity nested as random intercepts (1 | Block/Plant).

In both cases, posterior distributions were estimated from 4 parallel chains, each with 10,000 iterations (2,000 warmup), yielding 32,000 post-warmup draws. Model convergence was assessed using the potential scale reduction factor (*R̂* < 1.01) and effective sample size (ESS) for both bulk and tail. Divergent transitions and maximum treedepth warnings were monitored. *Post hoc* contrasts and estimated marginal means were computed using the ‘emmeans’ package version 2.0.3 with highest posterior density (HPD) intervals at 95% probability. Results are reported as posterior median differences (Δ̃) for viral load and as posterior median log-odds for symptoms severity, in both cases with 95% HPD credible intervals. Effects were considered significant when the 95% HPD interval excluded zero. For comparison with conventional frequentist inference, equivalent two-sided *P*-values (*P_eq_*) were obtained from the Bayesian probability of direction *PD* = max[*P*(Δ̃ > 0|data), *P*(Δ̃ < 0|data)] as *P_eq_* = 2(1 - *PD*). Joint tests of model terms are reported as χ^2^ statistics with associated *P*-values derived from the posterior distributions.

## 3. Results

### 3.1 Viral accumulation

Viral loads were quantified by absolute RT-qPCR at 14 dpi and are expressed as log_10_ viral genomes per ng of total RNA. ALSV infection was confirmed by RT-PCR, but viral RNA accumulated at levels close to the detection threshold and was therefore not analyzed quantitatively. BLLM were fitted separately for each quantified virus (TRV, TuMV and ArLV1) with Arabidopsis genotype × Infection type as fixed effects and Block and Biological replicate (nested within Block) as random intercepts.

#### 3.1.1 TRV accumulation

The BLMM revealed significant effects of Arabidopsis genotype (χ^2^ = 70.140, 7 d.f., *P* < 0.001), Infection type (χ^2^ = 8.370, 2 d.f., *P* = 0.015), but not their interaction (χ^2^ = 18.606, 14 d.f., *P* = 0.181) on TRV RNA1 accumulation. Averaged across host genotypes, ALSV coinfection significantly reduced TRV accumulation relative to single infection (Δ̃ = −0.203 log_10_, 95% HPD = [−0.338, −0.060], *P_eq_* = 0.004), whereas ArLV1 coinfection did not (Δ̃ = −0.126 [−0.267, 0.016], *P_eq_* = 0.089).

At the genotype level, ALSV coinfection strongly reduced TRV accumulation in *dcl2 dcl4* (Δ̃ = −0.389 [−0.782, −0.005], *P_eq_* = 0.050), *ein2-1* (Δ̃ = −0.511 [−0.857, −0.150], *P_eq_* = 0.005), and *jin1* (Δ̃ = −0.382 [−0.771, −0.003], *P_eq_* = 0.051) relative to single TRV infection, while ArLV1 coinfection had significant effect only in *dcl2 dcl4* (Δ̃ = −0.441 [−0.820, −0.043], *P_eq_* = 0.026) (Fig. 1A). Among host genotypes, *drm1 drm2*, *ein2-1*, *jin1*, *npr1-1*, and *sid2-1* all accumulated significantly more TRV than wild-type when averaged across coinfection treatments (in all cases *P_eq_* ≤ 0.004).

**FIGURE 1.**
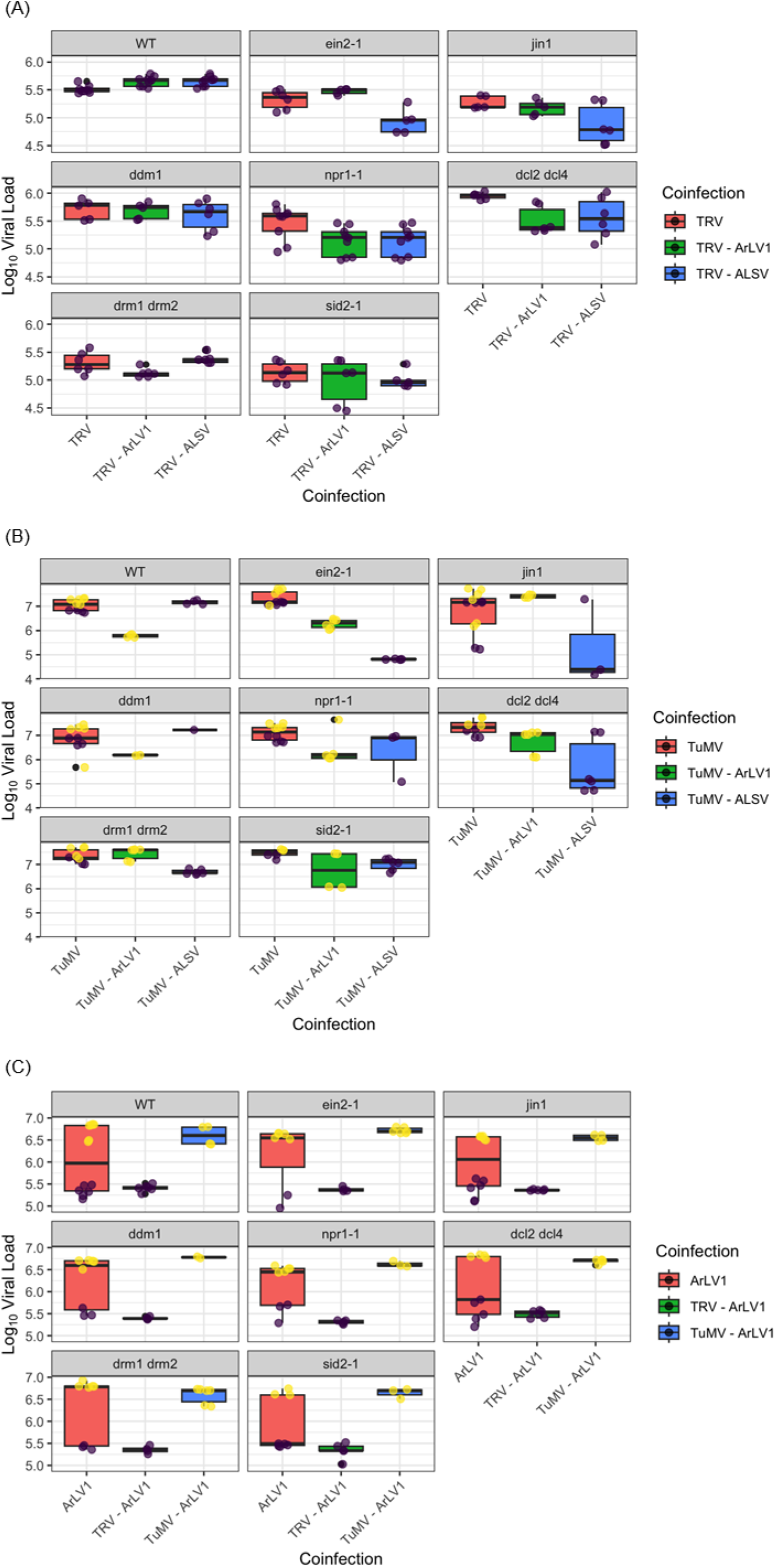
Viral accumulation (log_10_ viral genomes per ng of total RNA) at 14 dpi in *A. thaliana* Col-0 wild-type and defense-pathway mutants. (A) TRV accumulation in single and coinfections with ArLV1 and ALSV. (B) TuMV accumulation in single and coinfections with ArLV1 and ALSV. (C) ArLV1 in single and coinfection with TRV and TuMV. Horizontal lines represent the posterior medians and boxes the 95% HPD credible intervals from the BLMM. Dots correspond to individual biological replicates (3 pools of 4 plants) and are colored according to the experimental Block.

#### 3.1.2 TuMV accumulation

The BLMM showed significant effects of Arabidopsis genotype (χ^2^ = 24.052, 7 d.f., *P* = 0.001), Infection type (χ^2^ = 28.112, 2 d.f., *P* < 0.001), and their interaction (χ^2^ = 3.366, 14 d.f., *P* < 0.001), indicating that the impact of coinfection on TuMV accumulation depended on host genotype. Averaged across plant genotypes, both ASLV (Δ̃ = −0.654 [−1.056, −0.305], *P_eq_* < 0.001) and ArLV1 (Δ̃ = −0.687 [−1.016, −0.269], *P_eq_* = 0.003) have a significantly negative effect in TuMV accumulation.

ALSV coinfection significantly reduced TuMV in *dcl2 dcl4* (Δ̃ = −1.693 [−2.353, −0.597], *P_eq_* = 0.002) and *ein2-1* (Δ̃ = −2.529 [−3.300, −1.421], *P_eq_* < 0.001), while ArLV1 coinfection significantly reduced TuMV accumulation in wild-type (Δ̃ = −1.236 [−2.294, −0.429], *P_eq_* = 0.009) and *ein2-1* (Δ̃ = −1.375 [−2.032, −0.406], *P_eq_* = 0.001) (Fig. 1B). Among host genotypes, *ein2-1* accumulated significantly less TuMV than wild-type when averaged across coinfection treatments (Δ̃ = −0.662 [−1.101, −0.012], *P_eq_* = 0.023).

#### 3.1.3 ArLV1 accumulation

The BLMM revealed no significant effects of Arabidopsis genotype (χ^2^ = 6.629, 7 d.f., *P* = 0.468), Infection type (χ^2^ = 0.364, 2 d.f., *P* = 0.834), or their interaction (χ^2^ = 9.254, 14 d.f., *P* = 0.815) on ArLV1 RNA2 accumulation.

### 3.2 Disease severity progression

Disease severity progression was modeled using ordinal BGLMMs with a cumulative distribution and logit link, fitting separate models for the TRV and TuMV pathosystems. The fixed-effects structure included Arabidopsis genotype × Infection type × dpi, with Batch and Plant (repeated measures nested within Batch) as random intercepts. Contrasts are reported on the log_10_-odds ratio scale; negative values indicate reduced symptom severity relative to the reference. Fig. 2 shows the temporal dynamics of symptoms severity for each Infection type.

**FIGURE 2.**
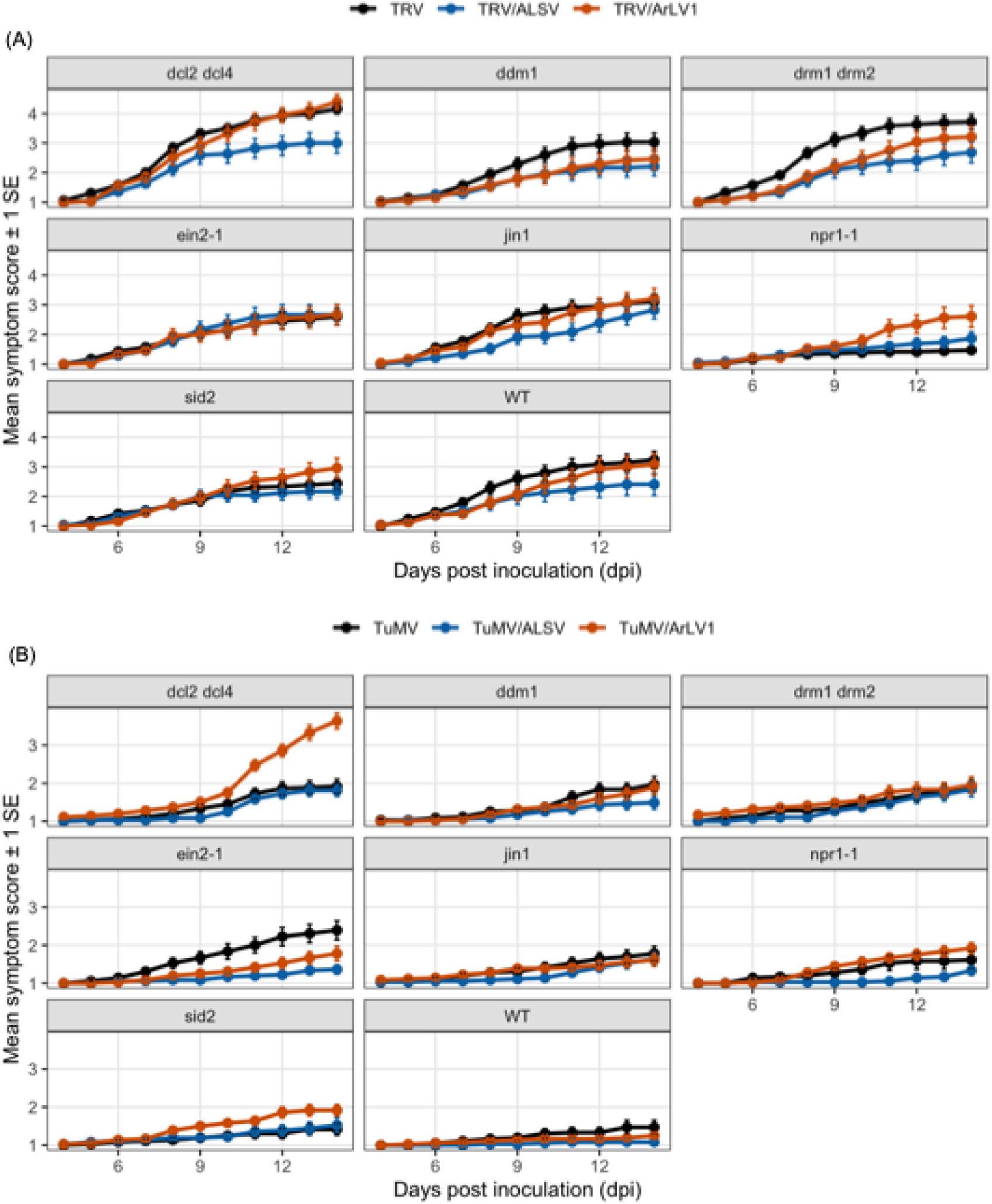
Temporal evolution of symptoms severity in single infections with acute viruses *vs* double infections with acute and latent viruses in different *A. thaliana* Col-0 genotypes. (A) TRV single infections (black), coinfected with ASLV (blue) or coinfected with ArLV1 (red). (B) TuMV single infections (black), coinfected with ASLV (blue) or coinfected with ArLV1 (red). Dots represent the mean symptoms severity and error bars ±1 SE.

#### 3.2.1 TRV pathosystem

The BGLMM analysis (7,260 observations, 658 plants) revealed a highly significant effect of Arabidopsis genotype (χ^2^ = 39.235, 7 d.f., *P* < 0.001) and Infection type (χ^2^ = 19.740, 2 d.f., *P* < 0.001), but not their interaction (χ^2^ = 18.004, 14 d.f., *P* = 0.206), on symptom severity progression (Fig. 2A). Averaged across genotypes, ALSV coinfection significantly attenuated TRV-induced disease severity relative to single TRV infection (posterior median log-odds = −3.800, 95% HPD = [−5.310, −1.926], *P_eq_* < 0.001). ArLV1 coinfection also tended to reduce severity, but this effect was not statistically significant (−1.240 [−2.980, 0.400], *P_eq_* = 0.150). The difference between ALSV and ArLV1 coinfection was significant (−2.540 [−4.540, −0.613], *P_eq_* = 0.011), indicating that ALSV was a more effective attenuator of TRV-induced disease.

Among host genotypes, *dcl2 dcl4* showed the highest symptom severity, significantly exceeding all other genotypes except *drm1 drm2*. In contrast, *npr1-1* showed the lowest severity, significantly lower than *dcl2 dcl4*, *jin1* and *sid2-1*. The temporal dynamics of symptom progression under single TRV infection and under TRV/ArLV1 and TRV/ALSV coinfections are shown in Fig. 2A.

#### 3.2.2 TuMV pathosystem

Also in this case, the BGLMM analysis (9,482 observations, 851 plants) found highly significant effects of Arabidopsis genotype (χ^2^ = 46.886, 7 d.f., *P* < 0.001), Infection type (χ^2^ = 38.608, 2 d.f., *P* < 0.001) and their interaction (χ^2^ = 30.884, 14 d.f., *P* = 0.006) (Fig. 2B). Averaged across genotypes, ALSV coinfection significantly reduced TuMV-induced disease severity (−4.140 [−6.380, −2.260], *P_eq_* < 0.001). In striking contrast, ArLV1 coinfection significantly increased disease severity (3.240 [1.580, 4.890], *P_eq_* < 0.001). The difference between the two latent viruses was large and highly significant (−7.400 [−9.460, −5.610], *P_eq_* < 0.001).

Among host genotypes, *dcl2 dcl4* again showed the highest severity, while wild-type showed the lowest (Fig. 2B). The Arabidopsis genotype × Infection type × dpi interaction revealed that ArLV1 coinfection increased severity in some genotypes (*e.g*., *drm1 drm2*, *jin1*, and *sid2-1*) while decreasing it in others (*e.g*., *ein2-1*, *npr1-1* and wild-type) (Fig. 2B), as reflected by the significant three-way interaction terms in the model.

Overall, ALSV consistently attenuated disease severity across both acute virus pathosystems, whereas ArLV1 produced genotype-dependent effects ranging from disease suppression to disease enhancement, particularly in the TuMV pathosystem.

## 4. Discussion

This work set out to determine whether and how latent viruses modulate the outcome of coinfection by an acute virulent virus, and whether these effects depend on the host’s immune context. By combining two latent *Secoviridae* with different levels of adaptation to *A. thaliana* (ALSV, not adapted; ArLV1, adapted), two acute viruses with distinct silencing-suppression strategies (TRV and its weak 16K VSR *vs* TuMV and its potent HC-Pro), and a panel of mutants affecting the main antiviral defense pathways, we show that coinfection outcomes are strongly genotype- and context-dependent. Using Bayesian hierarchical models we demonstrate that latent viruses modified both viral load and symptom dynamics, but the direction and magnitude of these effects varied with the viral partners and the host genotype.

Our viral-load results highlight the influence of RNA silencing and hormonal signaling pathways on viral accumulation, particularly in coinfection scenarios. As expected from the established role of DCL2 and DCL4 in antiviral RNA silencing (Bouché et al., 2006; Garcia-Ruiz et al., 2010), TRV showed a significant increase in accumulation levels in *dcl2 dcl4* plants. However, TuMV accumulation in single-infected *dcl2 dcl4* plants was undistinguishable from that in WT plants. The two acute viruses used here differ markedly in their VSRs, and this difference may contribute to the contrasted coinfection patterns observed (Martínez-Priego et al., 2008). It is also possible that due to its adaptation TuMV may bypass or suppress the DCL2/DCL4-dependent silencing pathway more efficiently.

Interestingly, coinfection with ALSV significantly reduced the accumulation of both acute viruses in dcl2 dcl4 plants, whereas coinfection with ArLV1 showed a more restricted effect. While TRV accumulation was reduced during TRV/ArLV1 coinfection compared to TRV alone, ArLV1 failed to alter TuMV accumulation in the same dcl2 dcl4 background. This contrast suggests that ALSV might trigger alternative or DCL2/DCL4-independent RNA silencing pathways capable of limiting acute viral accumulation broadly. Conversely, the inability of ALSV-like repression to occur during TuMV/ArLV1 coinfection could suggest an adaptive mechanism of ArLV1, reflecting its co-adaptation with the host to bypass or suppress host silencing pathways more efficiently.

Coinfection with ALSV consistently reduced the accumulation of both acute viruses in the ET-insensitive *ein2-1* mutant. This reduction of viral load under coinfection in *ein2-1*, observed both relative to wild-type and relative to single infection, implicates ET signaling in shaping coinfection outcomes.

The ordinal BGLMMs provided a nuanced view of disease severity progression complementing the viral-load findings. For the TRV pathosystem, ALSV coinfection significantly attenuated symptom severity, whereas ArLV1 produced a weaker, non-significant trend toward attenuation. For the TuMV pathosystem adapted to Arabidopsis, the contrast between the two latent viruses was stark: ALSV consistently reduced disease severity, while ArLV1 significantly increased it. This increase in TuMV-induced severity under ArLV1 coinfection is particularly noteworthy, as it represents a synergistic interaction where a latent virus enhances the pathogenicity of an acute partner. The ordinal model, by incorporating the full temporal dynamics of symptom progression (Arabidopsis genotype × Infection type × dpi), captured genotype-specific windows during which coinfection mattered, revealing that ArLV1 increased severity in some genotypes (*drm1 drm2*, *jin1* and *sid2-1*) while decreasing it in others (*ein2-1*, *npr1-1* and wild-type). These genotype-dependent effects underscore the importance of the host defense context in determining coinfection outcomes. Evaluating changes in disease progression between single and coinfections across mutant backgrounds, compared to the wild-type baseline, indicates that host defense pathways strongly modulate viral interactions. In particular, the SA and ET pathways appear essential for fine-tuning viral outcome.

Among these pathways, SA stands out as a major antiviral regulator. Its role well-documented in systems such as potato-potato virus Y interactions, where SA accumulation influences viral perception, defense activation, and symptom severity (Kutnjak et al., 2017). Similarly, in Arabidopsis, SA-dependent signaling contributes to resistance against viruses like cauliflower mosaic virus (CaMV) and CMV. However, the antiviral phenotypes of these mutants are far from uniform. In the CaMV pathosystem, loss-of-function mutants in the SA pathway (*eds5-1*, *NahG*, *pad4-1*, and *sid2-2*) do not exhibit enhanced susceptibility, unlike constitutive SA-signaling mutants (*cpr1-1* and *cpr5-2*), which show increased resistance (Love et al., 2007). Conversely, ET signaling mutants (*ein2-1* and *etr1*) display reduced susceptibility, as ET-dependent responses paradoxically enhance CaMV susceptibility and systemic spread (Love et al., 2007). These findings underscore the complexity of hormone-mediated defense, where SA often plays a central role, while JA and ET modulate outcomes through intricate crosstalk rather than a linear hierarchy (Takahashi et al., 2002; Tsuda et al., 2009; Sharma et al., 2024).

The variability of these effects across pathosystems further highlights the need for caution in interpreting mutant phenotypes. For example, SA and ET are essential for RCY1-mediated resistance to CMV-Y, whereas JA-insensitive mutations like *coi1* and *jar1* do not compromise this resistance (Takahashi et al., 2002; Takahashi et al., 2004). Similarly, *npr1* mutants retain resistance to certain strains of CMV and turnip crinkle virus, suggesting that defense signaling can branch below SA and sometimes operate independently of NPR1 (Love et al., 2005). Even *jin1*, typically associated with enhanced SA-dependent defenses, behaves unexpectedly as a highly permissive genotype in TuMV infections (Navarro et al., 2022). Thus, hormone-signaling mutants should be viewed as probes to explore interconnected defense sectors rather than as predictors of fixed antiviral outcomes. Their effects are context-dependent, shaped by the specific virus-host interaction, and reflect the dynamic interplay between hormonal pathways in plant immunity.

Even with a potent VSR, TuMV was not able to counteract the ALSV effect in the silencing defective mutant. This suggests that some hormonal pathways (ET, SA…) also play a role in the interaction between acute and latent viruses. One possible interpretation is that the stronger TuMV HC-Pro renders the acute virus more sensitive to changes in the silencing environment created by the latent partner and the host genotype, whereas the weak suppression by TRV 16K makes TRV accumulation more dependent on direct hormone-mediated defenses. This interpretation remains speculative and warrants direct experimental tests, for example through VSR-swap or complementation assays. ArLV1, adapted to Arabidopsis, maintains stable accumulation across all genotypes, suggesting it can bypass or attenuate host defenses, potentially facilitating the replication of acute viruses like TuMV through synergistic interactions. ALSV, poorly adapted to Arabidopsis, consistently shows very low viral loads appears to trigger host defense responses that restrict itself. However, its presence in coinfection with the acute viruses (TRV or TuMV) correlates with reduced accumulation of the acute partner in specific mutants (*dcl2 dcl4* and *ein2-1*). This suggests that ALSV may activate or enhance host defense pathways (*e.g*., RNA silencing or hormonal responses), which in turn restrict the acute virus. The persistence of ALSV low load even in defense-compromised mutants implies that residual mechanisms (*e.g*., DCL1 or AGO1) are sufficient to limit it, while still influencing the acute virus’s accumulation. Thus, while ArLV1 stable accumulation reflects its ability to evade host defenses and potentially aid acute viruses, ALSV low accumulation and impact on acute virus levels suggest it activates defense responses that restrict both itself and its coinfecting partner.

The Bayesian ordinal model uncovered that TuMV/ArLV1 was the only combination in which coinfection increased disease severity, and the only one in which the effect was strongly genotype-dependent, increasing severity in some genotypes (*drm1 drm2*, *jin1* and *sid2-1*) while decreasing it in others (*ein2-1*, *npr1-1* and wild-type). In contrast, ALSV consistently attenuated disease severity across both pathosystems. A plausible interpretation is that ArLV1, being adapted to Arabidopsis, engages host defenses more subtly, so that its impact depends more strongly on the genetic context, whereas the non-adapted ALSV triggers a more generic, and therefore more reproducible, modulation of the coinfecting virus. This interpretation is consistent with the suggested by the viral load data.

Additionally, our temporal analysis revealed a notable delay in the host response to coinfection: while no significant differences in viral accumulation or symptom severity were observed between single and mixed infections during the first week, a clear divergence emerged during the second week for some genotype. This pattern suggests a time-dependent activation of host defenses in response to the latent viral state. However, our coinfections were simultaneous, whereas in nature, latent viruses typically pre-exist in the host. Sequential (superinfection) scenarios may produce different outcomes, as suggested by cross-protection theory (Paudel and Sanfaçon, 2018). To further explore this hypothesis, it would be valuable to investigate whether such delayed responses persist in plants where the latent virus has been vertically transmitted, potentially establishing a more stable latent state. Testing whether prior establishment of ArLV1 or ALSV modifies the outcome of a subsequent TuMV or TRV infection is a natural extension of this work.

Two limitations of this study deserve mention. First, ALSV RNA accumulated near the qPCR detection limit, precluding quantitative analysis of the latent partner in the ALSV combinations. Second, viral load was measured at a single time point (14 dpi), and batch and virus treatment were partially confounded for some combinations in the qPCR experiment, so the corresponding contrasts should be interpreted with caution.

In conclusion, our analyses show that latent viruses are hidden players capable of modulating plant disease outcomes in coinfection contexts, through mechanisms that depend on both the viral partners and the host genetic background. Accounting for the full virome (pathogenic and nonpathogenic) present in the environment may therefore be necessary to predict viral emergence and disease severity in both wild and cultivated plant populations.

## Author Contributions

Guillaume Lafforgue: conceptualization, formal analysis, funding acquisition, resources, writing – review and editing. Santiago F. Elena: conceptualization, formal analysis, funding acquisition, resources, writing – original draft.

## Acknowledgments

We thank Francisca de la Iglesia and Ismael Rodrigo for excellent technical support and the rest of the EvolSysVir lab for fruitful discussions. We also thank Dr. René A.A. van der Vlugt (WUR, The Netherlands) for providing ArLV1 infectious material. This work was supported by grant PID2025-169610NB-I00 funded by MCIU/AEI/10.13039/501100011033 and by “ERDF a way of making Europe” and grant CIPROM/2022/59 funded by Generalitat Valenciana to S.F.E. G.L. was supported by the Plant Health and Environment (SPE) department and benefited from the Incentive Device for International Mobility (DIMI) of INRAE.

## Data availability Statement

Raw data and R code are available at Zenodo.org under doi: 10.5281/zenodo.23389679.

